# CROCCP2 acts as a human-specific modifier of cilia dynamics and mTOR signalling to promote expansion of cortical progenitors

**DOI:** 10.1101/2020.06.17.142976

**Authors:** Roxane Van Heurck, Marta Wojno, Ikuo K. Suzuki, Fausto D. Velez-Bravo, Jérôme Bonnefont, Emir Erkol, Dan Truc Nguyen, Adèle Herpoel, Angéline Bilheu, Catherine Ledent, Pierre Vanderhaeghen

## Abstract

The primary cilum is a central component of signalling during neural development, from regional patterning to neuronal differentiation. Here we focus on CROCCP2, a hominid-specific gene duplicate from CROCC (Ciliary Rootlet Coiled Coil), also known as rootletin, that encodes the major protein component of the ciliary rootlet. We find that CROCCP2 is highly expressed in the human fetal brain and not in other primate species. CROCCP2 gain of function in the mouse embryonic cortex results in decreased ciliogenesis, increased mTOR signalling, and increased cell size of radial glial cells, leading to increased generation of intermediate/basal progenitors and increased neuronal output. CROCCP2 impacts cilia dynamics and neurogenesis by inhibition of the IFT20 ciliary trafficking protein. Our data identify a human-specific protein that drives cortical basal progenitor expansion through modulation of ciliary dynamics.

## Introduction

The primary cilium is a sensory organelle present in most cells, regulating developmental processes and signalling (Goetz and Anderson, 2010). In neural development, it has been shown to contribute to regional patterning, neurogenesis, neuronal migration, and differentiation (Louvi and Grove, 2011). Indeed so-called ciliopathies, diseases caused by mutations in genes important for ciliary dynamics or function, display a variety of neurodevelopmental abnormalities (Lee and Gleeson, 2011; Louvi and Grove, 2011). As no protein translation occurs in the cilia, cilia trafficking is a major controller of ciliary dynamics and function. Intraflagellar transport (IFT) proteins, responsible for trafficking towards (IFTB) or from the cilia (IFTA), which thereby promote the dynamic growth or resorption of the cilia (Nachury and Mick, 2019).

The disruption of cilia-related genes has led to diverse and sometimes opposite phenotypes (Foerster et al., 2017; Guo et al., 2015; Tong et al., 2014; Wilson et al., 2012), reflecting the pleiotropy of cilia depending on cell and developmental context, as well as non-ciliary functions of some of the IFT proteins, for instance for centrosomal functions. Several ciliary genes have been shown to be crucial for neurogenesis (Guo et al., 2015). Ciliary dynamics has also been linked to proper apico-basal polarity (Higginbotham et al., 2013), delamination of neural progenitors (Das and Storey, 2014; Wilsch-Bräuninger et al., 2012), and the regulation of asymmetric divisions (Paridaen et al., 2013). While loss of function of ciliary genes then often results in impaired neurogenesis, decrased ciliogenesis has also been linked surprisingly to the amplification of specific populations of neural progenitors, most strikingly basal progenitors in the developing cerebral cortex (Foerster et al., 2017; Wilson et al., 2012). In these cases the impact of cilia on neurogenesis is thought to result in part from the modification of cilia-dependent signalling pathways, including mTOR and SHH (Foerster et al., 2017; Tong et al., 2014; Wilson et al., 2012). Ciliary dynamics has also been shown to influence directly cell-cycle speed in a direct fashion. Ciliary resorption has been shown to be required for cell cycle reentry (Pruski et al., 2016; Spalluto et al., 2012), thereby influencing directly the balance of self-renewal vs. differentiation of neural progenitors (Kim et al., 2011; Li et al., 2011).

The mechanisms of neurogenesis are thought to be largely conserved among all vertebrates, but divergence in neurogenic patterns are thought to play a major role in the evolution of shape and size of the brain. This is most striking for the primate cerebral cortex, which has undergone massive increase in size and complexification in the human lineage (Amadio and Walsh, 2006; Lui et al., 2011; Rakic, 2009). Cortical progenitors constitute a diverse set of cells located in the proliferative zones lining the lateral ventricles of the dorsal telencephalon, from which all cortical pyramidal neurons are generated (Kriegstein and Alvarez-Buylla, 2009; Dimou and Götz, 2014). The main cortical progenitors are radial glial cells (RGC), which constitute the ventricular zone (VZ) and divide at its apical surface. RGC undergo multiple rounds of asymmetric cell divisions, thereby enabling the generation of diverse types of neurons while maintaining a pool of progenitors. In primates, RGC display stemness for a much longer period (for several months for human RGC instead of days in the mouse), and thus generate more neurons throughout development (Van den Ameele et al., 2014; Astick and Vanderhaeghen, 2018). Species differences are also linked to other types of progenitors, which are more prominent in species with a larger cortex, and therefore thought to contribute in a crucial way to its evolutionary expansion. These are the intermediate/basal progenitor cells (IPC), and the outer/basal radial glial cells (oRGC), which both divide at more basal levels in the cortex, thereby contributing to generate additional neurogenic niches (Borrell and Götz, 2014; Lui et al., 2011; Sun and Hevner, 2014). The IPC lack any apico-basal processes and divide basally to create the subventricular zone (SVZ). The oRGC, particularly expanded in the human cortex, share epithelial features of RGC and divide basally to create the outer subventricular zone (oSVZ), and can undergo multiple rounds of self-renewing divisions, thus providing an important additional source of increased neuronal output.

Divergent neurogenic patterns between human and non-human primates are likely to be linked mostly to changes in regulatory networks controlling neural stem/progenitor cell self-renewal/differentiation balance (Kanton et al., 2019; Pollen et al., 2019). However recent work has uncovered the role in neurogenesis of “hominid-specific” (HS) genes, which result from recent gene duplications in hominid/human genomes (Dennis and Eichler, 2016). HS genes include >20 gene families that are highly and dynamically expressed during human corticogenesis, from early progenitors to maturing neurons (Charrier et al., 2012; Suzuki, 2020; Suzuki et al., 2018). These include human-specific NOTCH2NL genes, which increase the self-renewal potential of human cortical progenitors (Fiddes et al., 2018; Florio et al., 2018; Suzuki et al., 2018), and TBC1D3 and ARGHAP11B involved in basal progenitor amplification (Florio et al., 2015; Ju et al., 2016). However the function of the other HS genes, if any, remain unknown during neurogenesis.

Here we study CROCCP2, a HS gene duplicate highly expressed during human corticogenesis, that results from a partial duplication of the CROCC (Ciliary rootlet coiled coil) gene, also known as rootletin, a large coiled-coil protein that constitutes the ciliary rootlet (Yang et al., 2002). CROCC is so far mostly known to be required for maintenance of sensory cilia (Chen et al., 2015; Mohan et al., 2013; Styczynska-Soczka and Jarman, 2015; Yang et al., 2002, 2005), and has also been implicated in centrosome cohesion (Au et al., 2017; Conroy et al., 2012), but its function is otherwise largely unknown.

We show that CROCCP2 is highly expressed in human cortical progenitors, and CROCCP2 gain of function in the mouse cortex leads to amplification of basal progenitors together with decreased ciliogenesis. These effects are associated with increased mTOR signalling and require the interaction of CROCCP2 with IFT20, previously implicated in ciliary trafficking (Follit et al., 2006). Our data identify CROCCP2 as a human-specific gene involved in cortical neurogenesis and potentially link ciliary dynamics to human brain evolution.

## Results

### CROCCP2 emerged through partial duplication of CROCC/rootletin and is highly expressed during human corticogenesis

We previously identified > 20 gene families resulting from HS duplication and highly expressed during human corticogenesis, including paralogs of the CROCC gene (Suzuki et al., 2018). Among these CROCCP2 stood out as a gene dynamically expressed at a higher level than its ancestor gene CROCC.

Inspection of the human genome revealed the presence of seven CROCC-like duplicated loci including the ancestral CROCC and two pseudogenes (CROCCP2 and 3, refseq gene annotation). Only CROCC and CROCCP2 showed abundant expression during human corticogenesis (Figure 1). CROCCP2 is the result of a partial duplication of exon 13 to 21 of CROCC (Figure 1A). CROCC encodes a large protein (2017 AA) comprising two main domains, a head domain located at the N-terminus, followed by a large coiled coil repetitive domain (Yang et al., 2002) (Figure 1B). The CROCCP2 protein is predicted to be 111 AA long and corresponds to the middle part of the coiled coil domain (Figure 1B). Importantly, specific peptide sequences corresponding only to CROCCP2 (and not to CROCC or CROCCP3) have been identified in several human proteome studies, supporting its existence as a protein (Huttlin et al., 2017; Rolland et al., 2014).

**Figure 1.**
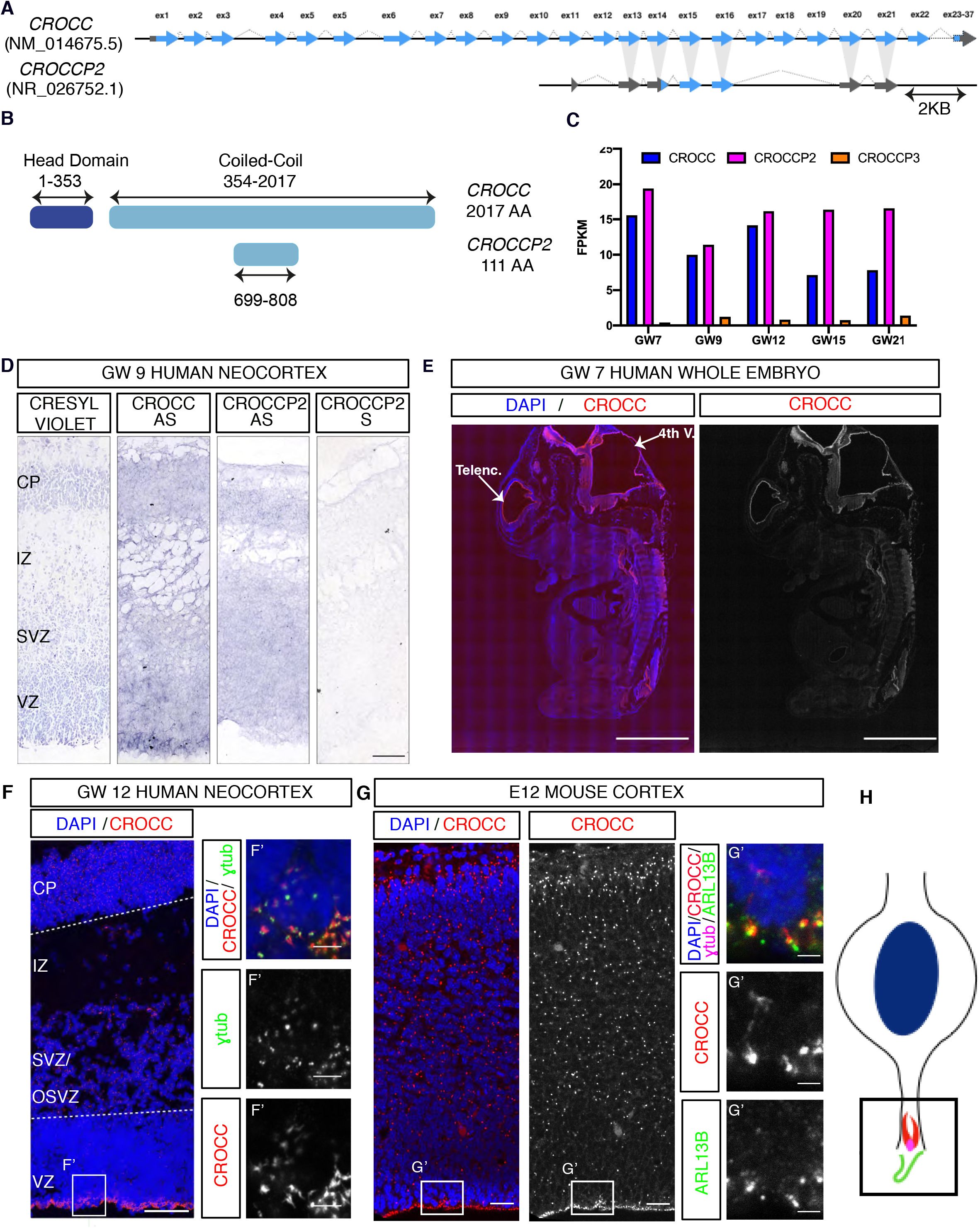
CROCCP2 is a HS duplicate highly expressed in human corticogenesis. (A) Schematic representation of CROCC and human-specific paralog CROCCP2 gene structure. Conserved exons are highlighted in light gray. Protein-coding and non-coding exons are depicted in blue and gray, respectively. CROCCP2 corresponds to duplication of exons 13 to 21 and is composed of 3 coding exons. (B) Putative protein structure of CROCC and CROCCP2. CROCC is composed of a small Head domain and a large Coiled-Coil repetitive domain. CROCCP2 corresponds to partial duplication of Coiled-Coil domain. (C) RNA-seq profile of CROCC HS gene family during human corticogenesis. (D) RNA in situ hybridization using specific probes for CROCC and CROCCP2 at 9 gestational weeks (GW). Note CROCC maximum intensity at apical surface of ventricular zone while CROCCP2 show diffuse expression through cortical wall. VZ, ventricular zone; SVZ, subventricular zone; IZ, intermediate zone; CP, cortical plate. Scale bar 100 υm. (E) CROCC protein expression on whole human embryo at GW7, note highest signal in telencephalic vesicles. scale bar 5 mm. (F) CROCC expression study during human corticogenesis (GW 12 cortical wall). F’-Inset at apical surface of ventricular zone, note elongated bundles located underneath basal body stained by φ tub. Scale bar 100 υm −5 υm in inset. (G) CROCC expression study during mouse corticogenesis (E14 cortex).(G’) Inset focusing on one primary cilia. Note localisation at the base of primary cilum. Scale bar 50 υm −3 υm in inset. (H) Shematic representation of interphase apical progenitor. See also Figure S1 & S2.

Inspection of primate and hominid available genomes revealed a diverse pattern of multiple CROCC-like coding sequences in the Catarrhini (the old world monkeys; Human, Chimpanzee, Gorilla, Orangutan, Gibbon, and Rhesus monkey), suggesting multiple species-specific genomic reorganizations. CROCCP2 locus is particularly divergent compared to the paralogous loci, but importantly the CROCCP2 protein studied here was uniquely found in human and chimpanzee. Inspection of the recently updated reference genome (panTro6) identified that, contrary to a single copy of CROCCP2 in human, Chimpanzee genome contains four tandemly duplicated regions encoding gene products similar to human CROCCP2. These data indicate that the CROCC gene was duplicated on multiple occasions in the Catarrhini lineages, probably because of the highly repetitive nature of its sequence and the tandem positions in a same chromosome, and leading to a gene duplicate encoding the CROCCP2 protein uniquely in human and chimpanzee.

To examine in detail the expression pattern of CROCCP2, we first relied on previously reported RNAseq datasets, in which analysis was tailored to distinguish with optimal sensitivity and specificity human-specific paralogs and their highly homologous ancestor genes (Suzuki et al., 2018). This revealed strong expression for CROCCP2 at higher levels than ancestor gene CROCC, and barely detactable levels of CROCCP3 (Figure 1C). Both genes displayed comparable but distinct patterns, with CROCC expression decreasing after 12 gestational weeks (GW) while CROCCP2 tends to maintain high expression throughout the examined stages. Moreover, analysis of recently released single transcriptome analyses of human, chimpanzee and macaque expression in fetal cortex and organoids (Nowakowski et al., 2017; Pollen et al., 2019) confirmed the expression of CROCCP2 in human fetal cortex, both in progenitors and neurons, while its expression was undetectable in the chimpanzee organoid-derived cortical cells (Figure S1). CROCCP2 is thus characterized by prominent and species-specific expression in the human fetal cortex.

To characterize the spatial expression of CROCCP2 we performed in situ hybridization on human fetal brain from 9 to 21 GW (Figure 1D, and data not shown). CROCC was found to be expressed throughout the cortical wall, with highest expression at the apical surface of the ventricular zone. In contrast CROCCP2 was expressed uniformly throughout the cortical wall (Figure 1D). We next examined CROCC protein localisation during human and mouse corticogenesis, taking advantage of a specific CROCC antibody. This revealed prominent expression throughout the central nervous system at early stages (Figure 1E). In the mouse and human developing cerebral cortex, CROCC immunoreactivity was found to be highest at the apical surface of RGC (Figure 1F-H), where it forms a large and elongated structure situated underneath the basal body (labeled with gamma-tubulin) and the primary cilium (labelled with ARL13B). A less intense, punctate labeling was also found throughout the cortical wall, consistent with centrosomal localization. To examine in more detail the expression of CROCC we turned to an in vitro 3D system of corticogenesis (Anja Hasche and P.V., unpublished data) enabling higher resolution imaging including STED microscopy. This revealed that CROCC formed large (up to several microns) striated substructures underneath the primary cilum (Figure S2A-C).

Examination of CROCC in cycling RGC in the mouse and human cortical progenitors also revealed that CROCC is present during mitosis at the level of the centrosomes (Figure S2D-G), sometimes in an asymmetric fashion, suggesting association with the mother centriole, as previously suggested during fly spermatogenesis (Chen et al., 2015).

### CROCCP2 gain of function in the mouse embryonic cerebral cortex increases the pool of basal progenitors

The expression data suggest that CROCC/CROCCP2 could play an important function during cortical neurogenesis. To test this we performed gain of function of CROCCP2 during mouse mid-corticogenesis (E13) using in utero electroporation of plasmids encoding tagged versions of human CROCCP2. CROCCP2 gain of function did not noticeably alter the overall cell distribution throughout the cortical wall (Figure 2A-C). However it led to an increase in the proportion of TBR2+ basal progenitors (Figure 2G-I), while the proportion of PAX6+ RGC or NEUROD2+ neurons remained unchanged (Figure 2D,E,F,J,K,L). Consistent with an amplification of basal progenitors, staining for mitotic marker pH3 revealed an increase in the proportion of mitotic cells located in basal compartments (abventricular mitoses) (Figure 2M-O) together with an increase in the number of total mitotic cells (Figure 2O). Almost all basally dividing cells were TBR2+ (97,92 % vs 100%; N= 9 controls / 12 CROCCP2 embryos), and very few SOX2+ (Median 0% for both control and CROCCP2 overexpression; N=9 CONTROL / 11 CROCCP2 embryos) (Figure S3A,B). These data suggest an increased generation and proliferation of IPC, which was further tested using EdU nuclear labeling to examine cell cycle dynamics. Indeed this revealed an increase in proliferation in VZ/SVZ (Figure 2P-R) together with increased proliferative index of TBR2+ cells, following CROCCP2 gain of function (Figure 2S), while SOX2+ RGC did not display such an increase (Figure S3C-E). Overall these data suggest that CROCCP2 overexpression leads to an increase in IPC that is mostly due to their increased conversion from RGC and increased proliferation.

**Figure 2.**
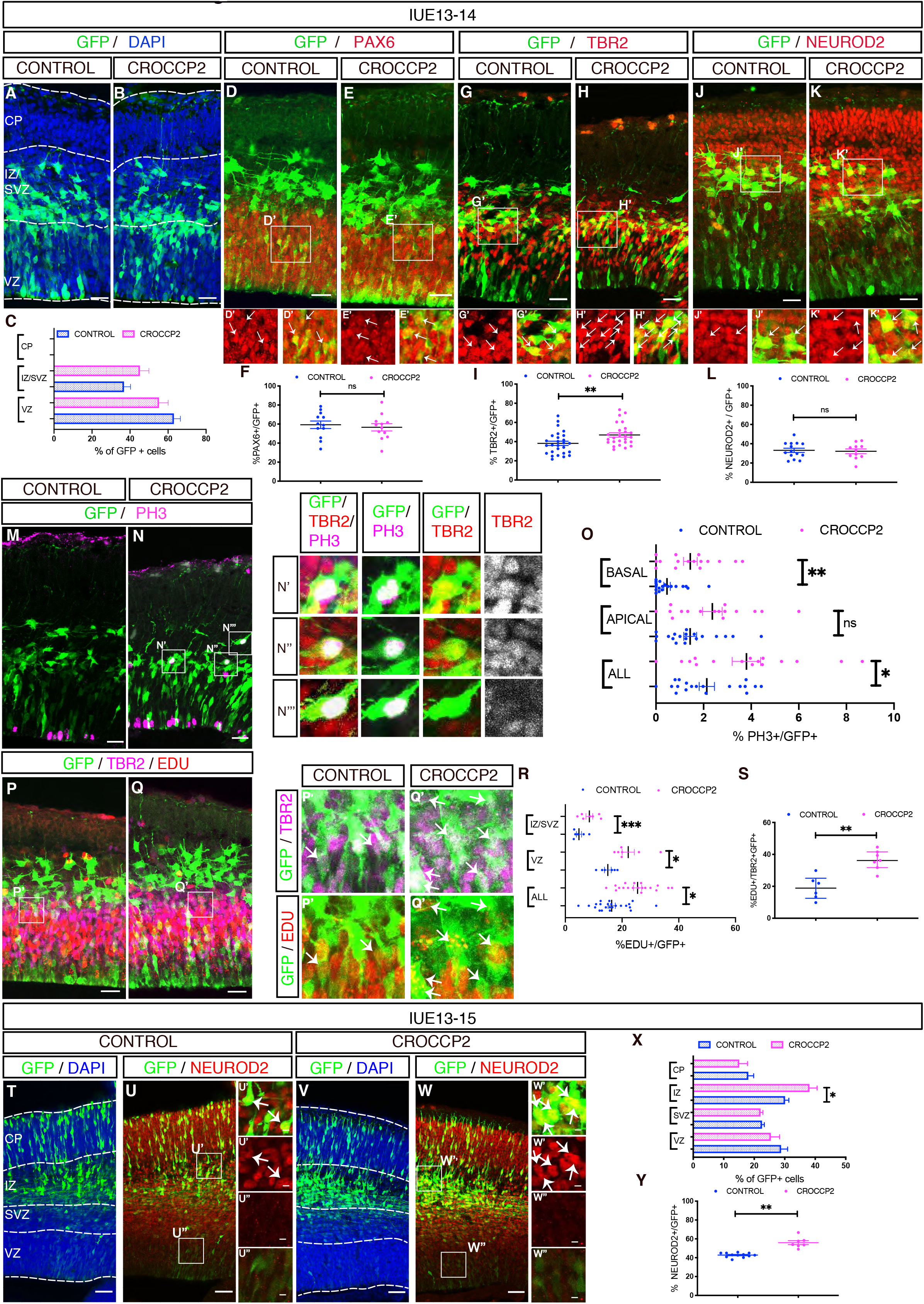
CROCCP2 gain of function leads to basal cortical progenitor amplification. In utero electroporation of pCIG (CONTROL) or Pcig-MYC-CROCCP2 (CROCCP2) at E13.5. (A,B) Immunofluorescence analysis of coronal sections of E14.5 brains stained for DAPI and GFP analyzing the distribution of transfected cells in CONTROL (A) and CROCCP2 (B) (C) Histograms showing the percentage of GFP+ cells in ventricular zone (VZ), subventricular zone and intermediate zone (SVZ/IZ) and cortical plate (CP) (*n*= 19 CONTROL embryos, 16 CROCCP2 embryos, *p* = 0,1977(VZ), *p* = 0,1631(SVZ), *p* =nd (CP). (D,E,G,H,J,K) Immunofluorescence analysis of coronal sections of E14.5 brains stained for GFP and PAX 6 (D,E) or GFP and TBR2 (G,H) or GFP and NEUROD2 (J,K) for CONTROL (D,G,J) & CROCCP2 (E,H,K). Arrows point to double positive cells. (F,I,L) Histogramms showing the percentage of GFP^+^ cells coexpressing either PAX6 (F), TBR2 (I) or Neurod2 (L). (n=12 CONTROL embryos, 26 CROCCP2 embryos, *p=0.6356* (PAX6); n=27 CONTROL embryos, 26 CROCCP2 embryos, \*\**p=0.0082* (TBR2), n=14 CONTROL embryos, 11 CROCCP2 embryos, *p*=0.787 (NEUROD2)). (M,N) Immunofluorescence analysis of coronal sections of E14.5 brains stained for GFP and phosphohistone H3 (PH3) in (M) CONTROL and (N) CROCCP2. (N’,N”,N’”) Inset showing GFP, PH3 and TBR2 immunofluorescence of basal dividing cells. Vast majority of cells are TBR2 positive. (O) Histogram showing the percentage and distribution of PH3^+^/GFP^+^ cells. (n=19 CONTROL embryos,16 CROCCP2 embryos \*\**p=0.0154* (ALL mitosis), *p= 0.0585* (APICAL), \*\**p<0.0102* (BASAL)). (P,Q) Immunofluorescence analysis of coronal sections of E14.5 brains stained for GFP, EDU and TBR2 in CONTROL (P) and CROCCP2(Q). (P’,Q’) Inset focusing on triple positive cells pointed by arrows. (R) Histogram showing the proportion and distribution of EDU^+^/GFP^+^ cells. (n=23 CONTROL embryos,19 CROCCP2 embryos *\*\*p=0.0367* (ALL cells), \*\**p=0.0183* (VZ), \*\**p<0.001* (SVZ/IZ)). (S) Histogram showing proportion of triple positive cells EDU^+^, TBR2^+^, GFP^+^. (n=6 CONTROL embryos, 7 CROCCP2 embryos, *\*\*p*=0,007). (A,B,D,E,G,H,J,K,M,N,P,Q) Scale bar 25 υm. (T-W) Immunofluorescence analysis of coronal sections of E15.5 brains stained for GFP and DAPI (T,V) or GFP and Neurod2 (U,W) in CONTROL (T,U) or CROCCP2 (V,W).Inset showing double positive (U’,W’) and GFP positive only (U’’,W’’) cells. scale bar 50υm (X) Histograms showing the percentage of GFP+ cells in ventricular zone (VZ), subventricular zone (SVZ), intermediate zone (IZ) and cortical plate (CP) (n = 16 CONTROL embryos,13 CROCCP2 embryos, *p* = 0.3422 (VZ), *p* = 0.6590 (SVZ), *\*p* = (IZ), *p* = 0.3511(CP)). (Y) Histogram showing proportion of GFP^+^ NEUROD2^+^ cells. (n=12 CONTROL embryos, 7 CROCCP2 embryos, \*\**p<0.0001)*. (C,F,I,L,O,R,S,X,Y) Data are represented as mean+/−SEM. Each dot represent an embryo, p values by Student t test. See also Figure S3.

We next examined the consequences of CROCCP2 gain of function at a later time point (48 h post electroporation). We observed an increased cell proportion in the intermediate zone (IZ) that contains newly generated neurons, together with an increased proportion of NeuroD2-positive neurons following CROCCP2 gain of function (Figure 2T-Y), while RGC and IPC appeared to be unchanged in number and proliferative behaviour (Figure S3F-Q). These data suggest that CROCCP2 gain of function leads to a transient increase in the expansion of IPC, followed by increased neuronal output, as expected given the known neurogenic committment of IPC (Martinez-Cerdeno et al., 2006).

### CROCCP2 decreases ciliary length of cortical progenitors

We next sought to determine the mechanism underlying CROCCP2-mediated increased generation of basal progenitors. Given the localization of CROCC at the level of the cilia, and the importance of this organelle in corticogenesis, we examined the impact of CROCCP2 on ciliary morphology in apical and basal progenitors. To this aim we performed co-electroporation of CONTROL/CROCCP2 expression vectors together with ciliary marker ARL13B fused with RFP expressing vector. This revealed that the primary cilia of RGC, typically located at the apical surface, were greatly shortened following CROCCP2 gain of function (Figure 3A-C), while there was no detectable effect on ciliary length in IPC or newborn neurons (Figure S4A-F). We further validated these data by examining endogenous ARL13B and acetylated tubulin on ex vivo cultures of mouse embryonic cortex, to maximize cellular resolution. This also revealed a striking shortening of the primary cilium in SOX2+ RGC and an increase in RGC devoid of any detectable cilium (Figure 3D-J).

**Figure 3.**
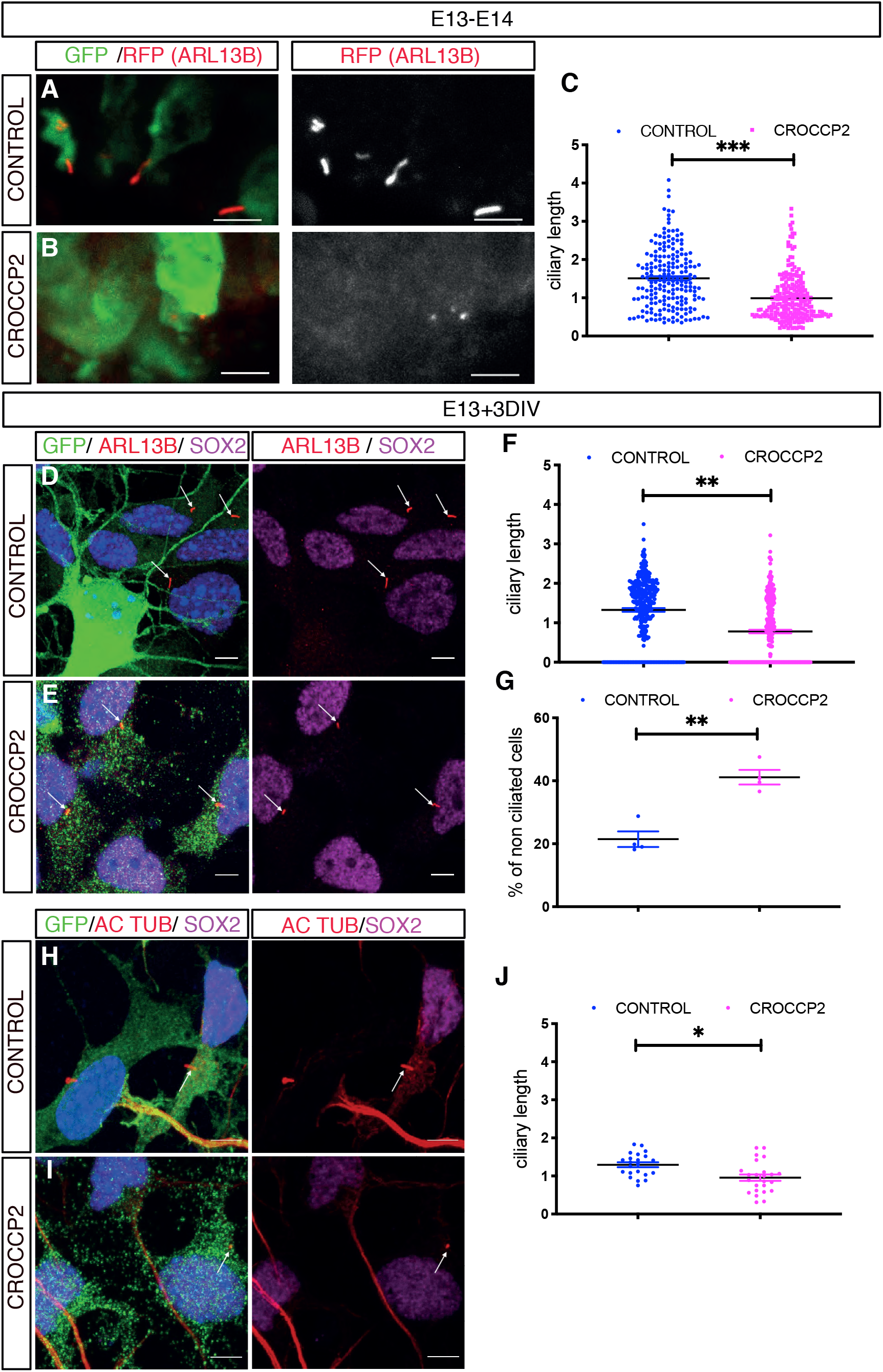
CROCCP2 gain of function leads to reduced ciliary length in apical cortical progenitors. (A,B) Mouse co in utero electroporation of ARL13B-RFP fused vector with pCIG (CONTROL)(A) or pCIG-MYC-CROCCP2 (CROCCP2)(B) at E13.5, followed by analysis at E14.5 through immunofluorescence staining of RFP and GFP. Scale bar 10 υm (A’,B’) Inset illustrates primary cilia at apical surface of ventricular zone. Scale bar 5υm. (C) Histogram showing primary cilia size at apical surface of venctricular zone (n= 210 cilias, 5 CONTROL embryos, 221 cilias, 4 CROCCP2 embryos \*\**p<0,001*) (D,E,H,I) Immunofluoresence of primary culture of mouse embryonic cortex infected with either pLenti-CIG-CONTROL (CONTROL) (D,H) or pLenti-CIG-MYC-CROCCP2 (CROCCP2) (E,I) expressing lentivirus at E13 followed by analysis 72h later. Scale bar 10 υm. (D,E) Immunostaining of DAPI, GFP, SOX2 and ARL13b. (H,I) Immunostaining of DAPI, GFP, acetylated tubulin (AC TUB) and SOX2. (F) Histogram showing ciliary length in SOX2^+^ cells using ARL 13B (n=4 experiments, 332 cilias CONTROL, 326 cilias CROCCP2*,** p<0.0001*) and (G) Histogram showing the percentage of non ciliated SOX2^+^ cells (n=4 experiments \*\**p=0.00286*). (J) Histogram showing ciliary length in SOX2^+^ cells using acetylated tubulin (n=2 experiments, 22 CONTROL cilias, 24 CROCCP2 cilias, \*\**p=0.0029*). (C,F,J) Data are presented as mean +/− SEM, one dot represent one cilia, p values by Student t test. (G) Data are presented as median +/− IQ, each dot represents mean percentage of non ciliated cells per experiment, p values by Mann-Whitney test. See also Figure S4.

Overall these data indicate that CROCCP2 overexpression leads to a strong reduction in ciliary size in mouse cortical progenitors.

### CROCCP2 interacts with IFT20 regulator of ciliary trafficking

We then sought to identify the molecular link(s) between CROCCP2 and primary cilia shortening. A previous proteomic study described direct interactions between CROCCP2 and IFT20 (Rolland et al., 2014), an IFTB protein that regulates the transit from Golgi to the base of the primary cilia (Follit et al., 2006; Olatz Pampliega et al, 2013). Co-transfection of CROCCP2 with IFT20 in HEK cells revealed their co-localization at a perinuclear localization (Figure 4 A,B) partially overlapping with Golgi marker GM130 (Figure 4C,D). This finding was confirmed in human cortical progenitors in vitro, in which overexpressed CROCCP2 was co-localized with endogenous IFT20 (Figure 4E,F). Similar Golgi-like localization was found in mouse cortical progenitors overexpressing CROCCP2 (Figure S5 A,B).

**Figure 4.**
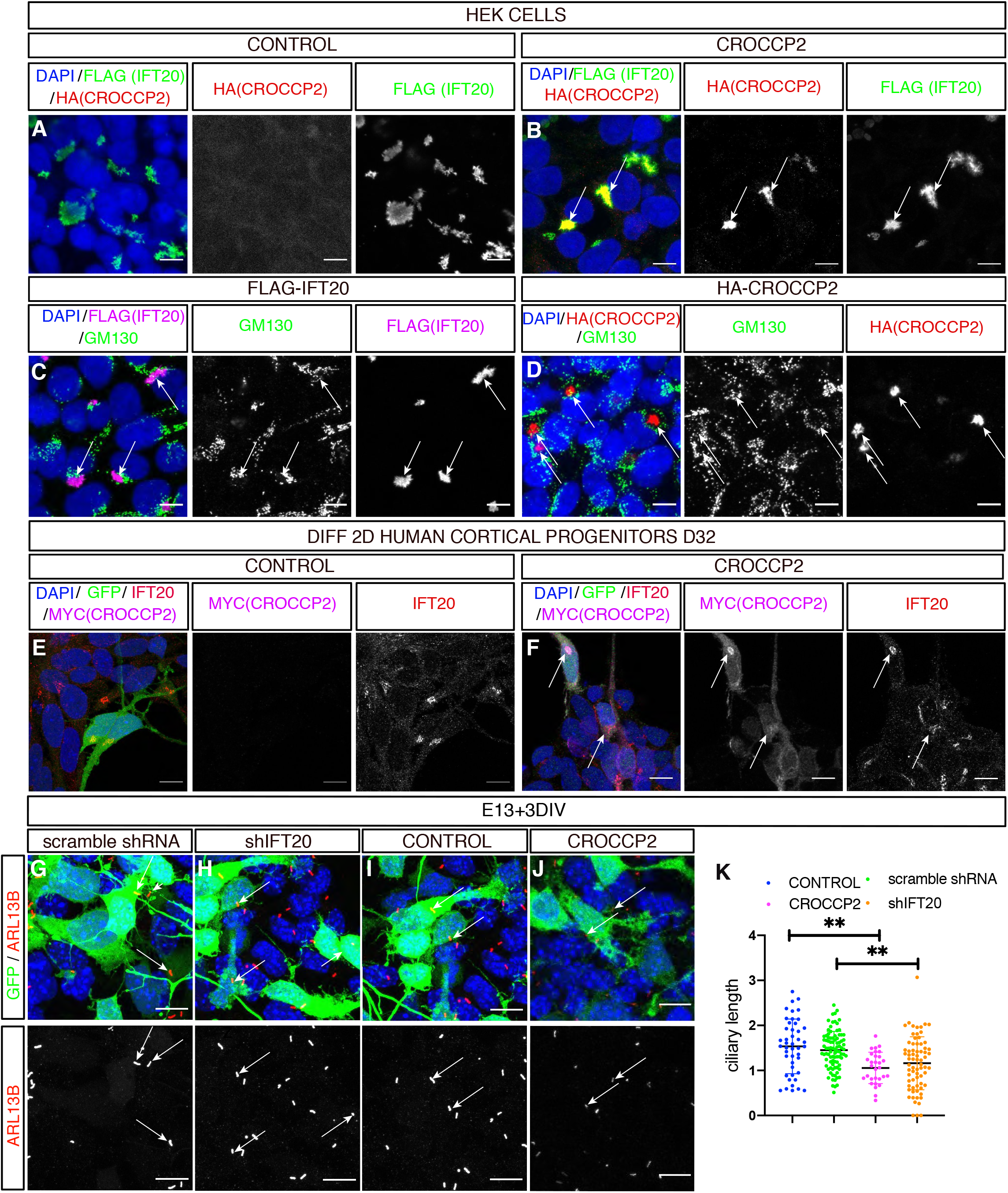
CROCCP2 co-localizes with IFT20 at the Golgi level. (A,B) Immunofluorescence staining of HEK cells co-transfected with pCIG-HA-CROCCP2 (HA (CROCCP2)(B) or pCIG (CONTROL)(A) & p3xFLAG-IFT20 (FLAG(IFT20)). Note co-localization of CROCCP2 & IFT20 tagged proteins in CROCCP2 expressing cells. (C,D) Immunofluorescence staining with GM130 on HEK cells transfected either with p3xFLAG-IFT20(C) or pCIG-HA-CROCCP2(D) expressing vector. Note partial co localization of IFT20 with GM130(C) and CROCCP2 with GM130(D). (E,F) Human 2D in vitro corticogenesis (D32), infected with pLenti-CIG-CONTROL(CONTROL)(E) or pLenti-CIG-MYC-CROCCP2(MYC(CROCCP2)(F) at D25 and analyzed 6 days after infection. Note Myc signal colocalization with IFT20 endogenous signal (F). (G,H,I,J) Immunofluorescence staining of DAPI, GFP and ARL13B on mouse cortex primary culture after ex utero at E13,5 using scramble shRNA (G), IFT20 shRNA (shIFT20) (H), pCIG (CONTROL)(I), PCIG-MYC-CROCCP2 (CROCCP2)(J) vectors. Data analyzed 72h after transfection. Arrows indicate short primary cilias. (K) Histogram showing ciliary length measured through ARL13b immunostainings. Data are represented as mean +/− SEM. Each dot represent one cilia. (n=2 experiments, 45 cilias CONTROL, 90 cilias scramble shRNA, 49 cilias CROCCP2, 72 cilias shIFT20, *p=0.8053* (CONTROL vs scramble shRNA), \*\**p= 0.0005* (CONTROL vs CROCCP2), \*\**p=0.0008* (CONTROL vs shIFT20), **p=0.0020 (scramble shRNA vs shIFT20), **p*=0.0015 (*scramble shRNA vs CROCCP2), p=0.7801 (CROCCP2 vs shIFT20). P values by one way ANOVA followed by post hoc Tukey test. (A,B,C,D,E,F,G,H,I,J) Scale bar 10 υm. See also Figure S5 and S6.

### CROCCP2 impacts cortical neurogenesis through IFT20 inhibition

Co-localization of CROCCP2 and IFT20 at the Golgi level suggest that CROCCP2 could impact ciliogenesis by inhibiting IFT20 trafficking function required for ciliogenesis.

To test this hypothesis we first examined the effect of IFT20 loss of function in the mouse embryonic cortex, using in utero electroporation of IFT20-targeting shRNAs (Figure S6). The knock-down of IFT20 led to the shortening of primary cilia in vitro, similar to the effect of CROCCP2 gain of function (Figure 4G-K). Moreover, the downregulation of IFT20 in vivo led to similar effects as CROCCP2 gain of function, as we observed an increase in the proportion of cells undergoing basal mitosis (Figure 5A,B,E) and IPC (Figure 5C,D,F).

**Figure 5.**
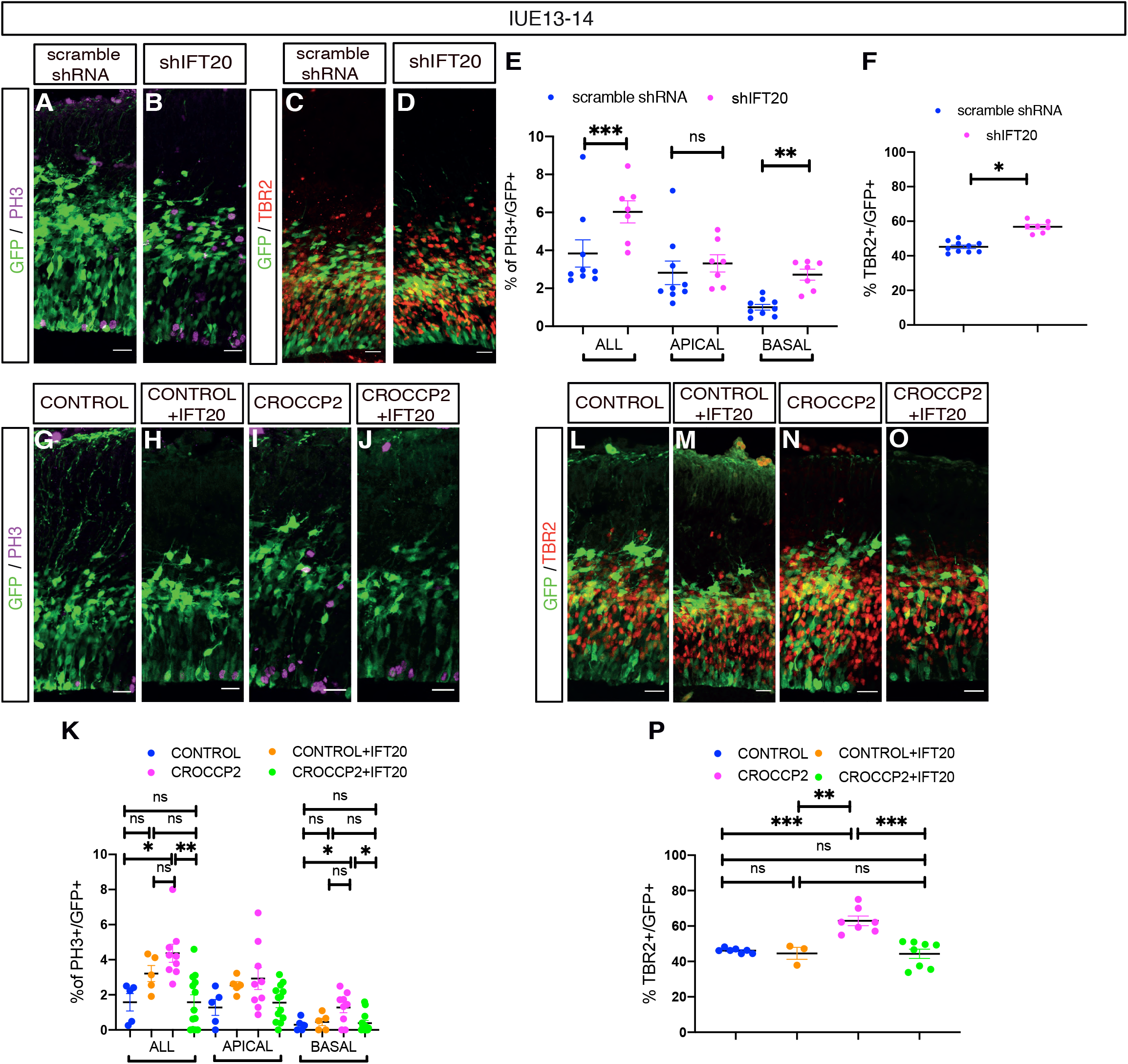
CROCCP2 affects cilia and neurogenesis through IFT20 downregulation. (A,B,C,D,G,H,I,J,,L,M,N,O) In utero electroporation of scramble shRNA, shRNA against IFT20 (shIFT20), pCIG (CONTROL), PCIG+p3XFLAG-IFT20 (CONTROL+IFT20), pCIG-MYC-CROCCP2 (CROCCP2), PCIG-MYC-CROCCP2+p3XFLAG-IFT20 (CROCCP2+IFT20) at E13.5. (A,B,G,H,I,J) Immunofluoresence of coronal sections of E14,5 brains stained with GFP & Phosphohistone H3 (PH3) or (C,D,L,M,N,O) stained with GPF & TBR2. Scale bar 25 υm. (E) Histogram showig the percentage of PH3^+^/GFP^+^ cells and their localisation. (n=9 scramble shRNA embryos,7 shIFT20 embryos, \**p=0.0399* (ALL mitosis), *p= 0.05482* (APICAL), \*\*\**p<0.0001* (BASAL). (F) Quantification of TBR2^+^/GFP^+^ progenitors (n=10 scramble shRNA embryos, 7 shIFT20 embryos, \*\*\**p<.0001*) (K) Histogram showing the percentage of PH3^+^/GFP^+^ positive cell and their distribution. (n=5 CONTROL embryos, 5 CONTROL+IFT20 embryos, 9 CROCCP2 embryos, 11 CROCCP2+IFT20 embryos; ALL mitosis : *p=0,2828* (CONTROL vs CONTROL+IFT20), \*\**p= 0.0073* (CONTROL vs CROCCP2), *p*>0,9999 (CONTROL vs CROCCP2+IFT20*), p=0,4691* (CROCCP2 vs CONTROL+IFT20), *p=0,1506* (CONTROL+IFT20 vs CROCCP2+IFT20) \*\*\**p=0.0005* (CROCCP2 vs CROCCP2+IFT20). BASAL mitosis: *p=0,9773* (CONTROL vs CONTROL+IFT20), \**p=0,0452* (CONTROL vs CROCCP2), *p= 0.9918* (CONTROL vs CROCCP2+IFT20), *p=0,1179* (CONTROL+IFT20 vs CROCCP2), *p=0,9971* (CONTROL+IFT20 vs CROCCP2+IFT20), \**p=0.0159* (CROCCP2 vs CROCCP2+IFT20). (P) Histogram showing the percentage of TBR2^+^/ GFP^+^ (n=7 CONTROL embryos, 3 CONTROL+IFT20 embryos, 7 CROCCP2 embryos, 8 CROCCP2+IFT20 embryos : *p=0,9835* (CONTROL vs CONTROL+IFT20), \*\*\**p= 0.0002* (CONTROL vs CROCCP2), *p*>0,9409 (CONTROL vs CROCCP2+IFT20*), **p=0,0013* (CROCCP2 vs CONTROL+IFT20), *p=0,9999* (CONTROL+IFT20 vs CROCCP2+IFT20) \*\*\**p<0,0001* (CROCCP2 vs CROCCP2+IFT20). (E,F,K,P) Data are presented as mean +/− SEM, each dot represent an embryo. (E, F) P values by Student t test (K,P) P values by one way Anova & Tukey post hoc test.

These data indicate that IFT20 loss of function leads to decreased ciliary length and increased basal progenitor amplification, in a way that is strikingly similar to the gain of function of CROCCP2. This supports the hypothesis that CROCCP2 and IFT20 may interact functionally. We therefore tested whether IFT20 gain of function could block the effects of CROCCP2, by performing their combined gain of function in the mouse cortex (Figure 5G-P). This revealed that IFT20 gain of function could completely block the effects of CROCCP2 on basal mitoses (Figure 5G-K) and IPC expansion (Figure 5L-P), suggesting that the effects of CROCCP2 require the inhibition of IFT20.

### CROCCP2 gain of function leads to increased mTOR signaling in cortical progenitors

What could be the downstream mechanism linking CROCCP2, ciliary dynamics and neurogenesis? The primary cilium has been shown to regulate many signalling pathways, including a direct inhibition of the mTOR pathway (Boehlke et al., 2010; Foerster et al., 2017), which was also recently found to be hyperactivated selectively in human cortical cortical progenitors and not in other primate species (Pollen et al., 2019).

We therefore examined mTOR activity following CROCCP2 gain of function (Figure 6). We first looked at cell size focusing on the surface of the apical endfoot of RGC in interphase (Figure 6A-E), as it is known to be increased by mTOR (Foerster et al., 2017). This revealed that CROCCP2 gain of function in RGC leads to the increase of RGC apical endfoot surface (Figure 6A-C). We next examined the levels of phospho-S6 protein as a classical readout of the mTOR pathway activity, on cortical progenitors transfected with CROCCP2. This revealed an important increase in phospho-S6 levels following CROCCP2 gain of function on mouse cortical progenitors (Figure 6F-H). Overall these data indicate that CROCCP2 activates the mTOR pathway in RGC, which could constitute a species-specific molecular effector of subsequent basal progenitor generation and amplification.

**Figure 6.**
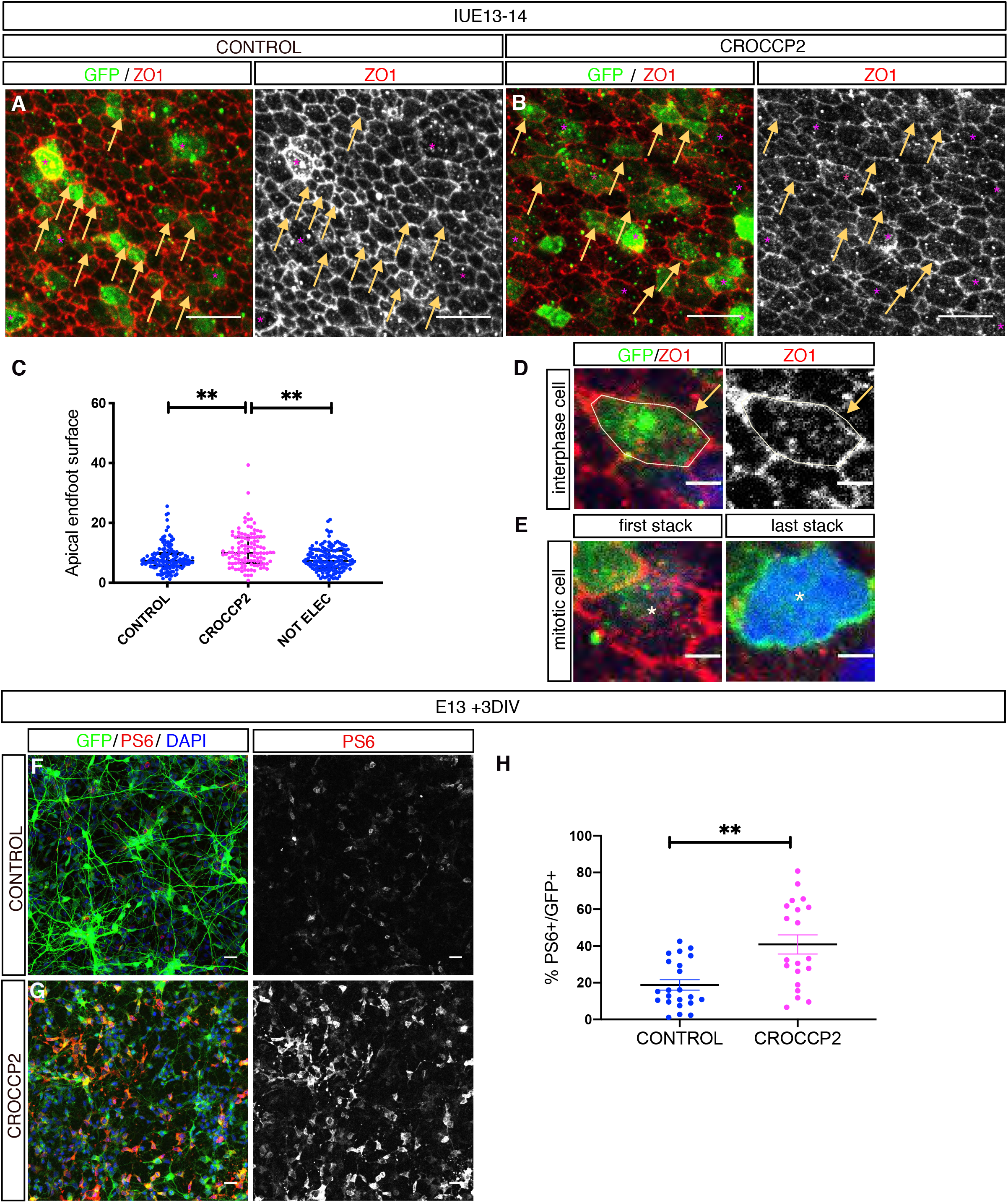
CROCCP2 gain of function leads to mTOR activation in cortical apical progenitors. Flat mount analysis of in utero electroporation using pCIG(CONTROL)(A) or pCIG-MYC-CROCCP2 (CROCCP2)(B) performed at E13.5 and fixed 24h later (A,B,D,E). Whole mount staining of en-face view of the ventricular zone using DAPI, GFP and ZO1 to stain apical endfoot. (C) Histogram showing surface measurement of apical endfoot. Data are represented as mean +/− SEM. One dot represent one apical endfoot. (n =136 apical endfoot CONTROL - 3 embryos,116 apical endfoot CROCCP2 - 4 embryos,152 apical endfoot of non electroporated (NOT ELEC) cortex sides -2 embryos, \*\**p<0.0001* (CONTROL vs CROCCP2), \*\**p<0.0001* (NOT ELEC vs CROCCP2), *p=0.8261* (NOT ELEC vs CONTROL). P values by one way Anova & Tukey post hoc test. (A,B)Scale bar 10 υm. Arrows point to interphase progenitors apical endfoot-stars mark mitotic progenitors. (D,E) Apical progenitors are classified in interphase (D) or mitotic (E) cells using dapi and extended Z stack imaging. Scale bar 2 υm. (F,G) Immunofluorescence analysis of mouse embryonic cortex primary culture infected with either plenti-CIG (CONTROL)(F) or plenty-CIG-MYC-CROCCP2 (CROCCP2)(G) at E13.5 and analyzed 72h later. DAPI, GFP and pS6 immunostainings as a readout of the MTOR pathway activity, scale bar 25 υm. (H) Histogram showing the percentage of pS6^+^/GFP^+^ cells. Data are represented as mean +/− SEM, 1 dot represent 1 microscopic field (n=4 experiments, 22 microscopic fields CONTROL,20 microscopic fields CROCCP2, 22631 cells CONTROL, 20382 cells CROCCP2) \*\**p=0.004* Student t test.

## DISCUSSION

Here we identify CROCCP2, a hominid-specific gene duplicate that is selectively expressed in human cortical progenitors. CROCCP2 negatively influences cilia dynamics and promotes the expansion of basal progenitors, at least in part through interaction with IFT20 ciliary trafficiking protein, together with increased mTOR signalling.

CROCCP2 emerged in evolution from a partial duplication of CROCC/rootletin. CROCC is required for the maintenance of specialized cilia of sensory cells such as photoreceptors, but its implication in non-specialized primary cilium remains unclear (Chen et al., 2015; Mohan et al., 2013; Yang et al., 2005). Intriguingly, CROCC was previously proposed to play a role in intracellular trafficking (Yang and Li, 2005). CROCC also functions to promote centrosomal cohesion by interacting with C-Nap1 (Bahe et al., 2005), the disruption of which leads to dwarfism and microcephaly (Floriot et al., 2015). These data suggest that CROCC may have an ancestral function in cortical neurogenesis, while CROCCP2 displays species-specific effects on cilia and IPC expansion. Future work should determine whether CROCC can function in link with CROCCP2 in the context of neurogenesis.

Indeed our data rather suggest that CROCCP2 acts on ciliary dynamics by interacting with IFT20 at the level of the Golgi apparatus. IFT20 is a well-known IFTB protein, further underlining the important role of primary cilia during corticogenesis. Interestingly, IFT20 was previously proposed to be required to epithelial-mesenchymal transition in kidney cells (Han et al., 2018), a processus mechanistically linked to delamination of apical progenitors and their conversion into basal progenitors (Itoh et al., 2013). It was also shown to be required for normal ciliogenesis and proliferation in adult neural stem cells in the mouse dentate gyrus (Amador-Arjona et al., 2011).

The primary cilium is involved in complex interplays with the mTOR pathway and autophagy (Boehlke et al., 2010; Pampliega and Cuervo, 2016). Hyperactivation of the mTOR pathway was previously linked to the absence of primary cilia (Boehlke et al., 2010; Foerster et al., 2017), and increased mTOR activity was proposed as a mechanism linking decreased ciliary dynamics and increased generation of basal progenitors (Foerster et al., 2017; Wilson et al., 2012), in direct line with our results. Interestingly, the mTOR pathway was shown recently to be hyperactivated in the human fetal cortex compared to macaque and chimpanzee, most strikingly in oRGC (Pollen et al., 2019). It will be interesting in this context to explore further the relationships between CROCCP2 and mTOR in human oRGC, despite the difficulty to study them experimentally, especially given the many implications of this signalling pathway in brain diseases (Lipton and Sahin, 2014). Other potential signalling pathways should also be explored that could link ciliary function and neurogenesis, such as SHH and Wnt (Kostic et al., 2019; Park et al., 2018), but also YAP, recently linking apical centrosome anchoring in RGC and IPC generation (Shao et al., 2020).

CROCCP2 appears to exert effects primarily on RGC, but the main result of its gain of function in the mouse is amplification of IPC basal progenitors. Human brain evolution is caracterized by an expansion in basal progenitor compartiments, associated with increased brain size, increased neuronal number and increased brain folding (Lui et al., 2011). While oRGC have been the center of recent focus on human neurogenesis given their amplification in the human cortex, IPC are likely to be important for human cortical size and its evolutionary expansion. Indeed, impairement of IPC generation and amplification by Tbr2 gene dirsuption leads to severe microcephaly in humans (Baala et al., 2007), and amplification of IPC has been proposed to be a key feature of increased cortical size in amniotes (Cárdenas et al., 2018). Moreover we find that IPC themselves appear to be influenced by CROCCP2: as they display higher proliferation rates even without cilia dynamics alteration, this could reflect indirect effects on IPC generation, or additional more direct effects of CROCCP2 on basal progenitors.

So far a handful of HS gene duplicates have been shown to impact on cortical progenitors, ARHGAP11B, NOTCH2NLB/C, and TBC1D3 (Fiddes et al., 2018; Florio et al., 2015; Ju et al., 2016; Suzuki et al., 2018), but many more are expressed actively during human corticogenesis (Florio et al., 2018; Suzuki et al., 2018). Our data identify CROCCP2 as a new HS candidate with a functional impact of cortical evolution, further pointing to recent gene duplications as important molecular substrates for the increase in cortical size characterizing the human species.

## Acknowledgments

The authors thank members of the Vanderhaeghen lab for helpful discussions and advice and J.-M. Vanderwinden of the ULB Light Microscopy Facility for his support with imaging. A subset of the images were acquired on a Zeiss LSM 880 – Airyscan system (Cell and Tissue Imaging Cluster, CIC), supported by Hercules AKUL/15/37_GOH1816N and FWO G.0929.15 to Pieter Vanden Berghe, University of Leuven. This work was funded by the European Research Council (ERC Adv Grant GENDEVOCORTEX), the FWO, the FRS/FNRS, the AXA Research Fund, the FMRE/GSKE, the Fondation ULB (to P.V.). R.V. was supported by a predoctoral Fellowship of the FRS/FNRS, I.K.S. was supported by postdoctoral Fellowship of the FRS/FNRS, M.W. was supported by a predoctoral Fellowship of the FRIA.

## Author Contributions

Conceptualization and Methodology, R.V. and P.V.; Investigation, R.V., M.W., I.K.S., F.D.V.B., J.B., E.K., D.T.N., A.H., A.B., C.L. ; Formal Analysis, R.V., I.S., M.W., and P.V.; Key reagents, I.K.S., R.V.; Writing - Original Draft, R.V. and P.V.; Writing - Review & Editing, R.V., I.K.S., J.B., and P.V.; Funding acquisition, P.V.; Resources, P.V.; Supervision: C.L., P.V.

## STAR*METHODS

Detailed methods are provided in the online version of this paper and include the following :

- KEY RESSOURCES TABLES
- CONTACT FOR REAGENT AND RESOUCE SHARING
- EXPERIMENTAL MODEL AND SUBJECT DETAILS

- Human fetal tissue collection and preparation
- Mice

- In utero electroporation
- Cell lines
- METHOD DETAILS

- DNA constructs
- RNA sequencing and transcriptome analysis
- *In situ* hybridization
- Cortical differentiation of human ESC
- Lentiviral preparation
- Cell cycling labelling assay
- Mouse cortex primary culture
- En face ventricular zone whole mount
- Immunofluorescence staining
- Confocal microscopy
- Quantification and statistical analysis

- Statistical analysis

## CONTACT FOR REAGENT AND RESOURCE SHARING

Further information and requests for resources and reagents should be directed to the Lead Contact Pierre Vanderhaeghen.

## EXPERIMENTAL MODEL AND SUBJECT DETAILS

### Human fetal tissue collection and preparation

The study was approved by three relevant Ethics Committees (Erasme Hospital, Université Libre de Bruxelles, and Belgian National Fund for Scientific Research FRS/FNRS) on research involving human subjects. Written informed consent was given by the parents in each case.

Human fetuses were obtained following medical pregnancy termination. Fetuses aged 7 gestational weeks (GW), (9 GW, 12 GW, 15 GW, and 21 GW) were used for RNA sequencing and *in situ* hybridization of cortical tissue. All cases were examined with standard feto-pathological procedures and none displayed clinical or neuropathological evidence of brain malformation. As soon as possible after expulsion (less than 6 hours), the brain was removed using standard fetal autopsy procedure, frozen in liquid nitrogen for RNA extraction and embedded as a whole in OCT compound (Tissue-Tek Sakura), then snap-frozen in a 2-methylbutane on dry-ice bath for histological studies.

### Mice

All mouse experiments were performed with the approval of the Université Libre de Bruxelles Committee for animal welfare. Mouse housing, breeding and experimental handling were performed according to the ethical guidelines of the Belgian Ministry of Agriculture in agreement with European community Laboratory Animal Care and Use Regulations. Animals were housed under standard conditions (12 h light:12 h dark cycles) with food and water *ad libitum*. For *in utero* electroporation experiments, timed-pregnant mice were obtained by mating adult RjOrl:SWISS CD1 mice (Janvier, France). The plug date was defined as embryonic day E0.5 and the day of birth as P0.

### *In utero* electroporation

*In utero* electroporation was performed as previously described (Bonnefont et al 2019). Briefly, timed-pregnant mice were anesthesized with a ketamine/xylazine mixture at E13.5, and each uterus was exposed under sterile conditions. Plasmid solutions containing 1-2 mg/ml of DNA were injected into the lateral ventricles of the embryos using a heat-pulled capillary. Electroporation was performed using tweezers electrodes (Nepa Gene) connected to a BTX830 electroporator (five pulses of 25 V for 100 ms with an interval of 1s). Embryos were placed back into the abdominal cavity, and mice were sutured and placed on a heating plate until recovery. Embryos were collected 24 or 48 hours after *in utero* electroporation and perfused transcardiacally with ice-cold 4% paraformaldehyde. Brains were dissected and soaked in 4% paraformaldehyde overnight at 4 °C. They were then washed in PBS for 24 h then soaked in 30% sucrose solution overnight for cryopreservation. Brains were included in Tissue-TEK OCT compound (Sakura) at −20 °C overnight then stored at −80 °C. After equilibration the blocs were sectioned with 20-μm thickness with a cryostat (Leica).

### Cell lines

Human H9 embryonic stem cells (WiCell) were routinely propagated on mitotically inactivated mouse embryonic fibroblasts in Knockout DMEM (Thermo Fisher Scientific) supplemented with 20% Knockout Serum Replacement (Thermo Fisher Scientific), 1X Non-essential Amino Acids (Thermo Fisher Scientific), 1X Penicillin/Streptomycin (Thermo Fisher Scientific), 1X 2-Mercaptoethanol (Merck), 2mM L-glutamine (Thermo Fisher Scientific).

## METHOD DETAILS

### DNA constructs

Coding sequence of CROCCP2 was amplified by PCR using Prime Star DNA polymerase (TAKARA) from GW9 human fetal cortex cDNA. Primers were designed on the basis of the sequence of the GRCh38/hg38 reference human genome. The PCR fragment was then subcloned into the *pCAG::myc-IRES-eGFP* expression plasmid (Dimidschstein et al., 2013; Tiberi et al., 2012) by In Fusion cloning (Clontech). DNA fragment of *pCAG::myc-tagged CROCCP2-IRES-eGFP* was then amplified and cloned into lentiviral plasmid backbone (gift from Cecile Charrier) using In Fusion cloning. A *pCAG::3xHA-tagged CROCCP2-IRES-eGFP* expression vector and lentiviral expression vector were also generated. All constructs were verified by DNA sequencing and western blot. The *Flag-IFT20* sequence obtained from addgene plasmid (Cat#118033) was cloned into *pCAG::IRES-RFP* expression plasmid. The *RFP-tagged ARL 13 b* vector is a kind gift of Pr Heiko Lickert (Kinzel et al., 2010). Primers used to create the constructs are summarized in table S2.

The *IFT 20* shRNA sequences (5’-CCAAAGAAGCAGAGAACGA-3’) (5’-TTTATTGACCAATTTATA-3’) were cloned downstream of the *U6* promoter into the pSilencer2.1-CAG-Venus (pSCV2)-plasmid as previously described. Their efficiency was validated following immunostaining of primary culture of mouse cortical progenitors performed after *ex utero electroporation* using shRNA targetting mouse IFT20.

### Transcriptome analyses

Bulk RNAseq analysis was as described in (Suzuki et al., 2018). Publicly available primary and organoid samples from Pollen et al. 2019 were downloaded from GSE124299. CPM normalized cells were scaled after top 2000 variable genes were selected. Principle component analysis were performed and first 20 principal components were selected for non-linear dimensional reduction with t-stochastic neighbour embedding. Plotting tools of Scanpy were used for visual purposes (Wolf et al., 2018).

### *In situ* hybridization

*In situ* hybridization was performed using digoxigenin-labeled RNA probes (DIG RNA labeling kit, Roche) and alkaline phosphatase revelation (NBT/BCIP kit #SK-5400; Vector) using PCR-amplified or above-mentioned plasmid templates as previously described. Sense probes were used as a negative control for each gene tested. Images were acquired with a Zeiss Axioplan 2 microscope and a Spot RT3 camera using the Spot 5.2 software. The probes used in this study are summarized in table S3.

### Cortical differentiation of human ESC

Cortical differentiation from human ESC was performed using previously described protocol (Espuny-camacho et al., 2013) with slight modifications. At differentiation day -8, the hES cells were dissociated using PluriSTEM Dispase-II (Merck)/ Collagenase (Thermo Fisher Scientific) solution and plated on Growth factor-reduced matrigel (Corning) in Essential 8 Flex medium (Thermo Fisher Scientific). At differentiation day -2, the cells were dissociated using Stem-Pro Accutase (Thermo Fisher Scientific) and plated at low confluency (5,000–10,000 cells/cm^2^) on hES qualified matrigel (Corning) using Essential 8 Flex medium supplemented with 10 μM ROCK inhibitor (Y-27632; Merck). At differentiation day 0, the medium was changed to DDM (Gaspard et al., 2008) supplemented with B27 (Thermo Fisher Scientific) and 100 ng/mL Noggin (R&D systems), and the medium was replenished every day. After 16 days of differentiation, the medium was changed to DDM, supplemented with B27 (DDM/B27), and changed every day. At day 25, the progenitors were dissociated using Accutase and frozen using mFreSR (Stemcell technologies). Cortical progenitors were thawed on matrigel-coated coverslips 12-well using DDM/B27 supplemented with 10 μM ROCK inhibitor and fixed after 6 days using 4% paraformaldehyde for 20 min at 4° or ice-cold methanol for 6 min and then rinsed 3 times with PBS before immunostaining.

### Lentiviral preparation

HEK293T cells were transfected by packaging plasmids, *psPAX2* (Addgene #12260) and *pMD2.G* (Addgene #12259), and the transfer vector using Xtreme gene 9 transfection reagent (Sigma-Aldrich). Two days after transfection, culture medium was collected and viral particles were enriched by filtration (Amicon Ultra-15 Centrifuge Filters, Merck). Titration of every batch of lentiviral preparation was estimated following transduction of HEK293T cells. The hES cell-derived cortical cells were infected by the lentiviral constructs at differentiation day 25. The culture medium was changed the next day and phenotypes were analyzed 6 days after infection for CROCC & CROCCP2 subcellular localisation. Mouse primary cortex cultures were infected on the day of infection and phenotypes were analyzed 72h later.

### Cell-cycle labeling assay

Cell-cycle kinetic differences were assessed by labelling cortical progenitor cells using a nucleotide analog 5-ethynyl-2’-deoxyuridine (EdU; Merck) *in vivo* following 150 □g EdU injection into the peritoneal cavity of pregnant mice 1 hour before the sacrifice of the embryos. Detection of EdU was performed using Click-iT EdU Alexa Fluor 555 Imaging Kit (Thermo Fisher Scientific).

### Mouse cortex Primary culture

Mouse embryonic dorsal cortices were dissected under binocular at E13.5 in ice-cold HBSS (Thermo Fisher) containing with 1% Penicillin/Streptomycin (Thermo Fisher), 0.1 M HEPES (Thermo Fisher). They were then dissociated in Trypsin (Thermo Fisher) supplemented with 25mM EDTA (Merck) and 50 mg DNASE I (Sigma-Aldrich) for 30 min at 37°. Tubes were shaken every 10 min. Cortices were centrifuged and neutralized twice with NeuroBasal medium supplemented with 1% Penicillin/Streptomycin, 1% Glutamax, 1% Sodium Pyruvate, 1x B27 with vitamin A, 1x N-2 supplement (all from Thermo Fisher) and 1 mM N-acetyl cysteine (Sigma-Aldrich). Cells were passed through 0.4 micron cell strainer (Falcon), and then counted. Cells were then infected with adequate lentivirus and plated at 400,000 cells/3.5 cm^2^ on hES qualified-matrigel (Corning)-coated coverslips (Menzel-Gläser). For shRNA experiments, brain cortices were electroporated *ex utero* before being isolated and dissected as described above. In all experiments cells were cultivated for 72 h in culture medium as described above with addition of 10ng/mL human b-FGF (Peprotech) and culture medium was changed everyday. The cells were then rinsed with PBS and fixed either with 4% paraformaldehyde for 20 min at 4 °C or using a Glyoxal solution as previously described (Richter et al., 2018) before being washed 3 times in PBS.

### En-face ventricular zone wholemount

Wholemount staining of en-face view of the ventricular zone was performed as described in (Mirzadeh et al., 2010). Briefly, 24h after *in utero* electroporation, mouse embryos were collected and perfused transcardiacally with ice-cold 4% paraformaldehyde. Electroporated cortices were then dissected to completely expose the lateral wall and flatened out with the ventricle side up. Cortices were soaked in 4% paraformaldehyde overnight and then rinsed 3 times in PBS before overnight incubation in RIMS solution (Yang et al., 2014).

### Immunofluorescence staining

The tissues were washed with PBS for 10 min, then with PBS/0.3% Triton X-100 for 30 min. They were then quenched in 4 mM glycine solution for 30 min, except for whole mount staings, then blocked in PBS containing 0.3% Triton X-100 and 3% horse serum, for at least 1 hour. The tissues were incubated overnight at 4°C with primary antibodies in blocking solution. After three washes in PBS/0.3% Triton X-100, slides were incubated during 2 hours at room temperature with secondary antibodies in PBS/0.3% Triton X-100. After washing in PBS/0.3% Triton X-100, the tissues were mounted on a slide glass with DAKO glycergel mounting medium (DAKO) complemented with the DABCO anti-fading agent.

### Confocal microscopy

Confocal imaging was performed on a LSM780NLO confocal system fitted on an Observer Z1 inverted microscope (Zeiss) equipped with a Chameleon Vision II 690-1064 nm multiphoton laser (Coherent Europe). Fluorochromes were separated by linear unmixing using ZEN 2012 software (Zeiss). *In utero* electroporation were imaged using a Plan Apochromat 20x/0,8 dry objective, *in vivo* cilia imaging was done using LD C Apochromat 40x/1.1 water immersion objective and *in vitro* cilia imaging was done using LD C Apochromat 63x/1.4 oil immersion objective (Zeiss). Wholemount cortices were flattened into fluorodish (Ibidi) and covered with glass coverslip before image acquisition using the 63x objective on 4-5 Z-stacks spaced at 0.3 μm to cover the whole tight-junction region.Images were then processed using the Fiji/ImageJ software.

Sted images were acquired using a DMi8 inverted microscope (Leica) equipped with STED with 592nm (CW), 660nm (CW) and 775nm (pulsed) depletion lasers. HC PL APO 100x/1.4 oil objective was used. Acquisition was done with LAS X and deconvolution using Huygens Professional Version 17.1. For Sted imaging, ABBERIOR secondary antibodies were used.

## QUANTIFICATION AND STATISTICAL ANALYSIS

### Statistical analysis

Results are shown as mean±standard error (S.E.M.) of at least three biologically independent experiments or median with interquartile range when mean of experiments is presented. Student’s unpaired *t*-test was used for two group comparisons. Analyses of multiple groups were performed by a one-way analysis of variance (ANOVA) followed by post hoc Tukey’s test. For all tests, a *P*-value inferior to 0.05 was taken as statistically significant.

## Supplemental files

Supplementary figure S1-S6

Table S1 Key ressources table related to STAR methods

Table S2 Primers for HIS related to STAR methods

Table S3 Primers for DNA construction related to STAR methods

**Figure S1 (Related to Figure 1).**
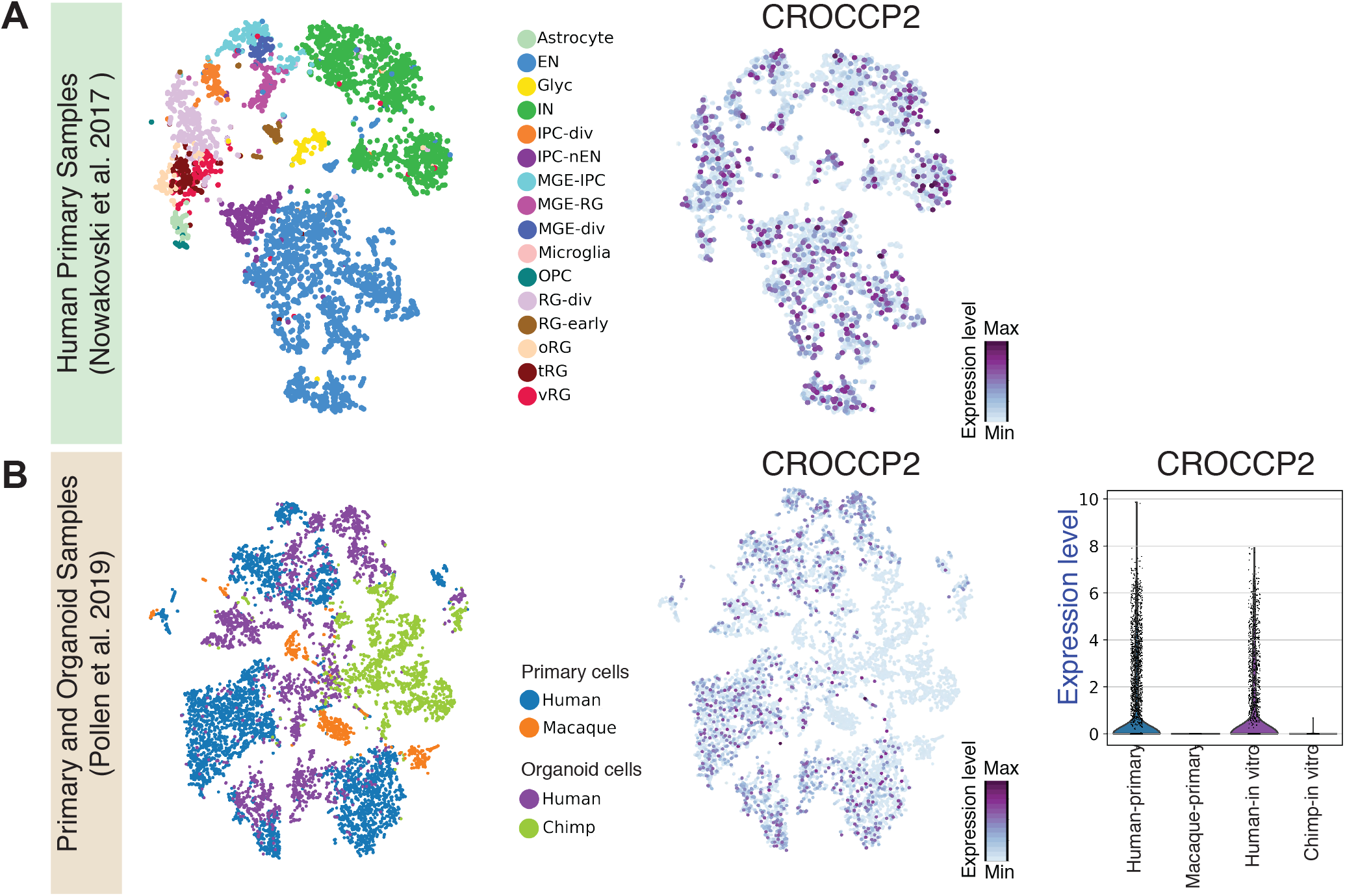
CROCCP2 expression in human cortical cells is species-specific. (A) Plotting human fetal cortex cells as provided from Nowakowski et al. 2017. Expression of CROCCP2 gene is colored (right). (B) t-stochastic neighbor embedding of top principal components of human and macaque primary samples and human and chimp organoid samples from Pollen et al. 2019. CROCCP2 expression is colored by expression level. Violin plots (right) shows CPM normalized expression of CROCCP2 gene for each sample. EN; excitatory neurons, Glyc; glycolysis, IN; interneuons, IPC-div; dividing intermediate progenitor cells, IPC-nEN; intermediate progenitors-newborn excitatiory neurons, MGE-IPC; medial ganglionic eminence-Intermediate progenitors, MGE-RG; medial ganglionic eminence-radial glia, MGE-div; dividing medial ganglionic eminence cells, OPC; oligodendrocyte precursor cells, oRG; outer radial glia, tRG; truncated radial glia, vRG; ventricular radial glia.

**Figure S2 (Related to Figure 1).**
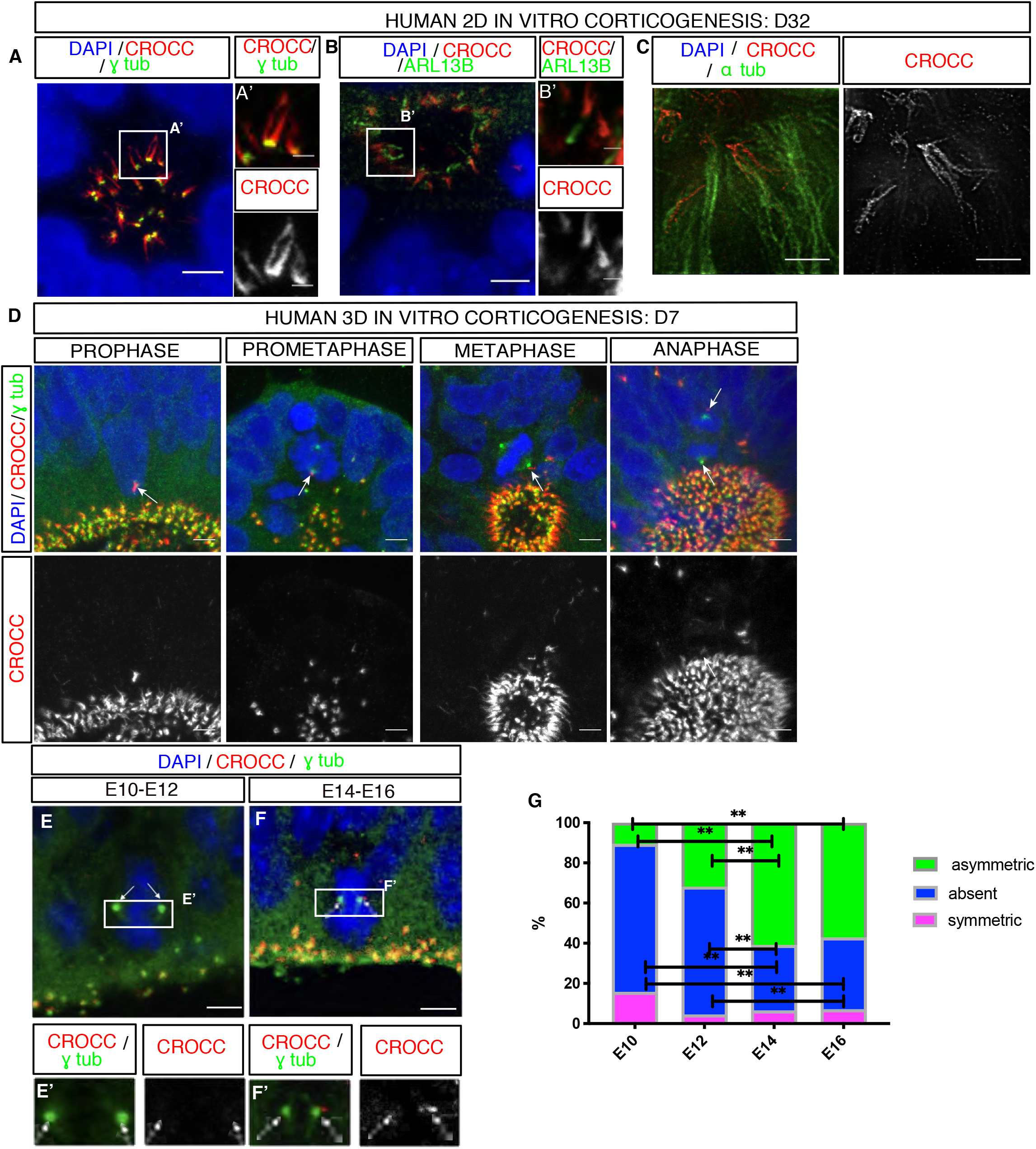
CROCC expression in cortical progenitors and displays dynamic expression during cell cycle. (A,B) Human in vitro corticogenesis depicting CROCC localisation at apical surface of cortical progenitors rosettes using both 2D (A) and 3D (B) in vitro differentiation model. (A) Co staining of DAPI, j tubulin & CROCC at apical surface of progenitors rossettes. CROCC is located just below basal body where it forms elongated structures.(A’) Inset showing high magnification (B) Co staining of DAPI, primary cilia (ARL 13B) and CROCC at apical surface of 3D neuroepithelial cysts. (B’) Inset showing high magnification. Scale bar 5 um & 1 um in inset. (C) Immunostaining of 2D human differentiation at D32 of CROCC and alpha tubulin imaged with STED at apical surface of progenitors rosettes. Note striated aspect of CROCC suggesting repetition of the same epitope. Scale bar 2 um. (D) Immunostaining of DAPI, CROCC and j tubulin focusing on mitotic apical progenitor at different stages of mitosis during human 3D in vitro corticogenesis. Note asymmetric localisation of CROCC at the centrosomes in prometaphase and metaphase. (E,F) Immunofluorescence staining of embryonic mouse brain coronal sections with DAPI, CROCC and j tubulin focusing on metaphase apical progenitors at different stages of mouse corticogenesis. Note absence of CROCC at the centrosomes during early stages of corticogenesis (symmetric mitosis) and asymmetric localisation of CROCC at the centrosomes at later stages of corticogenesis. (G) Histogram showing distribution of CROCC among the centrosomes at different stages of mouse corticogenesis. Note the switch from absent to asymmetric signal from E12 to E14. Data is represented as mean. (P value is 2 way ANOVA with Tukey post hoc. ABSENT : **p<0,001 (E10 vs E14),** p<0,001 (E10 vs E16), p=0,340 (E10 vs E12), **p <0,001 (E12 vs E14), *p=0,015 (E12 vs E16), p=0,458 (E16 vs E14). ASYMMETRIC **p<0,001 (E10 vs E14),** p<0,001 E14 vs E12), p=0,186 (E14 vs E16), **p<0,001 (E16 vs E10), p=0,093 (E12 vs E16), p=0,134 (E12 vs E10).

**Figure S3 (Related to Figure 2).**
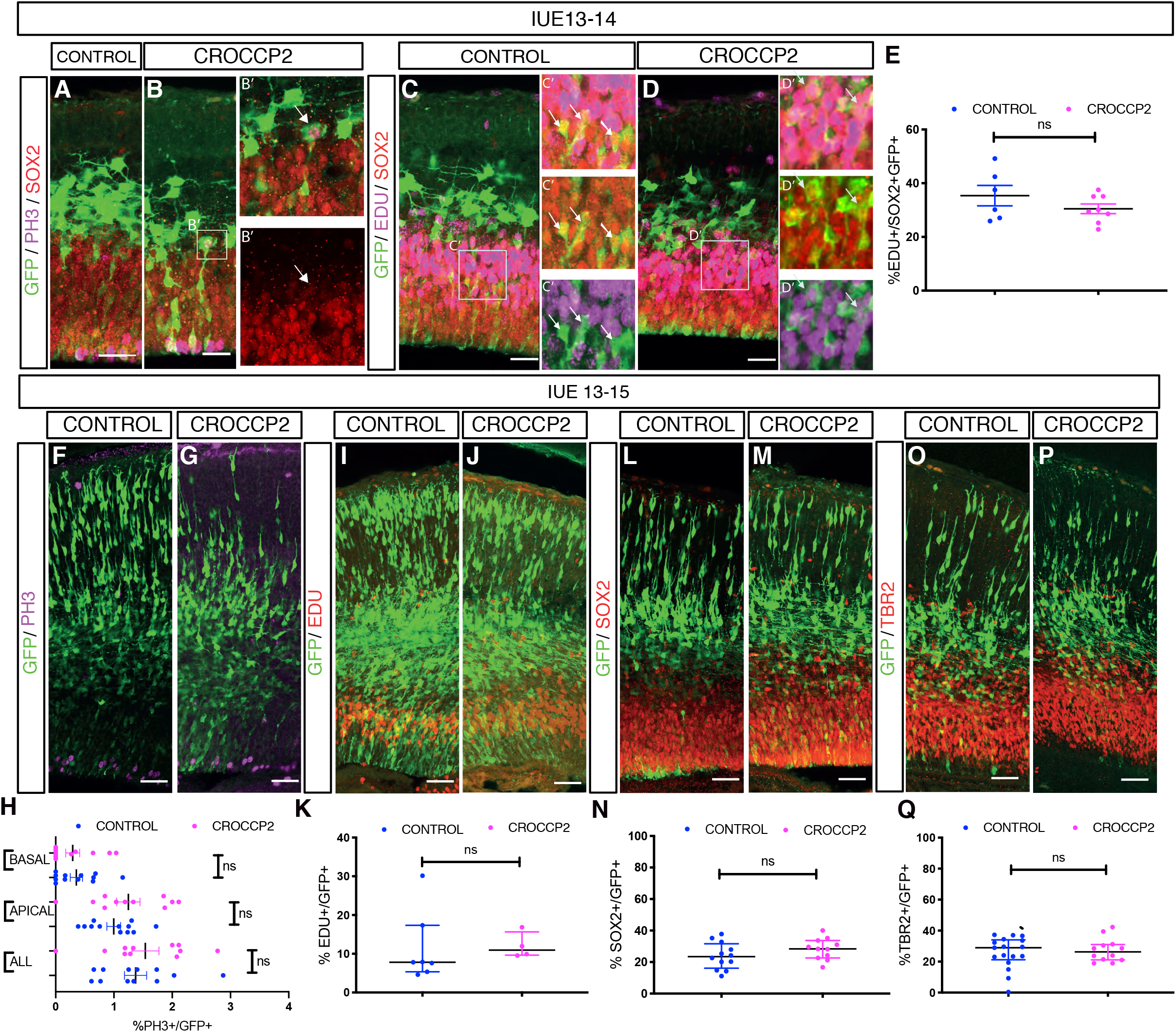
CROCCP2 overexpression does not increase the apical progenitor pool. (A,B,C,D,F,G,I,J,L,M,O,P) In utero electroporation of E13.5 mouse neocortex using pCIG (CONTROL) and pCIG-MYC-CROCCP2 (CROCCP2). (A,B) Immunofluorescence staining of coronal sections of E14.5 brains with GFP, SOX2 and PH3 in (A) CONTROL and (B) CROCCP2. (B’) Inset, note weakly positive SOX2 signal of basal dividing cells, (n=9 CONTROL embryos, 18 CROCCP2 embryos) (C,D) Immunofluorescence staining of coronal sections of E14.5 brains with GFP, EDU & SOX2 in CONTROL (C) and CROCCP2 (D),arrows points to GFP+/SOX2+/EDU+ cells. (A-D)scale bar 25 υm. (E) Histogram showing the percentageof EDU+/SOX2+ among GFP+cells (n=6 embryos each) p= 0,0631. Immunofluorescence staining of coronal sections of E15.5 brains with (F,G) GFP & PH3, (I,J) GFP & EDU (L,M) GFP & SOX2 (O,P) GFP & TBR2 in CONTROL (F,I,L,O) and CROCCP2 (G,J,M,P). scale bar 50 υm. (H) Histogram showing the quantification and distribution PH3+/GFP+. (n=11 CONTROL embryos,10 CROCCP2 embryos); p=0.577 (ALL mitosis), p= 0.2798 (APICAL), p=0.6963 (BASAL). (K) Histogram showing the percentage of EDU+/GFP+ cells (n=7 CONTROL embryos, 4 CROCCP2 embryos) p= 0,941 (N) Histogram showing the percentage of SOX2+/GFP+ (n=11 CONTROL emrbyos,10 CROCCP2 embryos) p=0.1866, (Q) Histogram showing the percentage of TBR2+/GFP+ (n=17 CONTROL embryos,12 CROCCP2 embryos) p=0.394. (E,H,K, N,Q) Data are represented as mean +/− SEM. Each dot represent an embryo. P values by Student’s t test.

**Figure S4 (Related to Figure 3).**
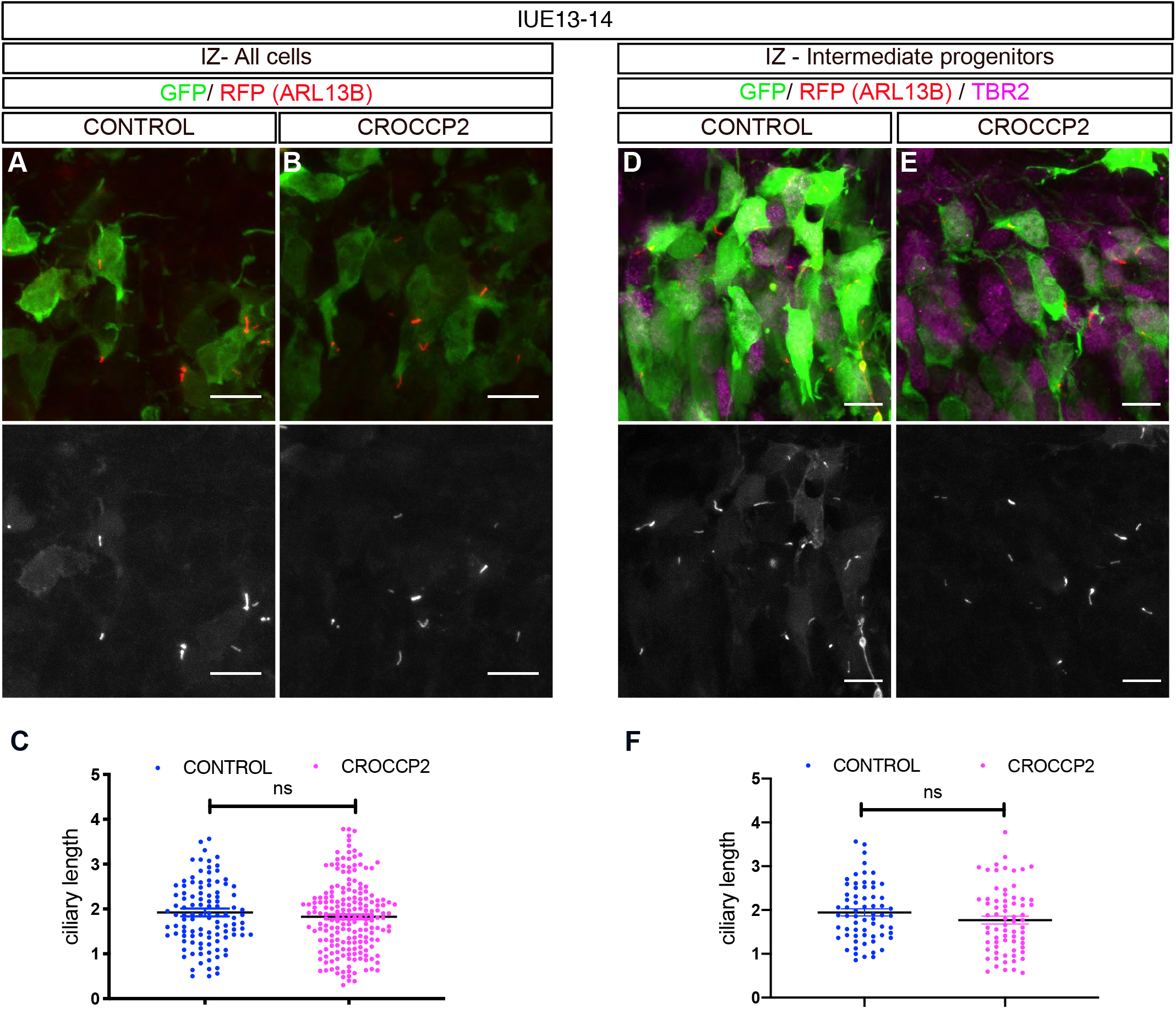
CROCCP2 gain of function does not alter ciliary length in basal progenitors or neurons. (A,B,D,E) Mouse co in utero electroporation of ARL13B-fused with RFP vector with Pcig (CONTROL) (A,D) or Pcig-MYC-CROCCP2 (CROCCP2) (B,E) at E13.5.(A,B,D,E) Immunofluorescence staining of mouse embryonic brain coronal sections at E14,5 focusing on the intermediate zone with (A,B) ARL13 labelled through RFP and GFP immunostaining or (D,E) RFP, GFP and TBR2 immunostaining. Scale bar 10 υm. (C) Histogram showing ciliary length in all GFP+ cells located in the IZ (n=108 cilias CONTROL - 5 embryos, n=193 cilia CROCCP2 - 4 embryos) p=0.339. (F) Histogram showing ciliary length in TBR2+/GFP+.(n=65 cilias CONTROL - 3 embryos, 70 cilias CROCCP2 - 3 embryos) p=0.1485). (C,F) Data are represented as mean +/− SEM, each dot represent one cilia, p values by Student’s t test.

**Figure S5 (Related to Figure 4).**
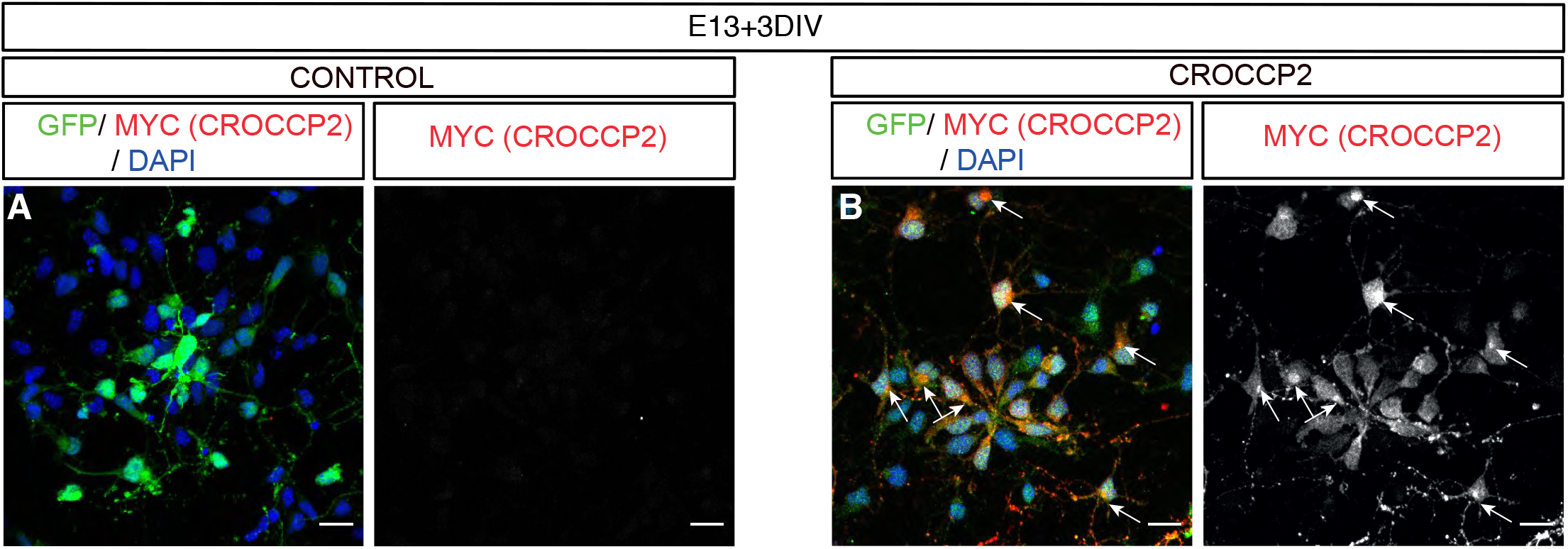
CROCCP2 shows golgi like subcellular localisation in mouse cortical progenitors. (A,B) Immunofluorescence analysis of mouse cortex primary culture stained with GFP, & MYC (CROCCP2). Note perinuclear staining of CROCCP2. Scale bar 20 υm.

**Figure S6 (Related to Figure 4).**
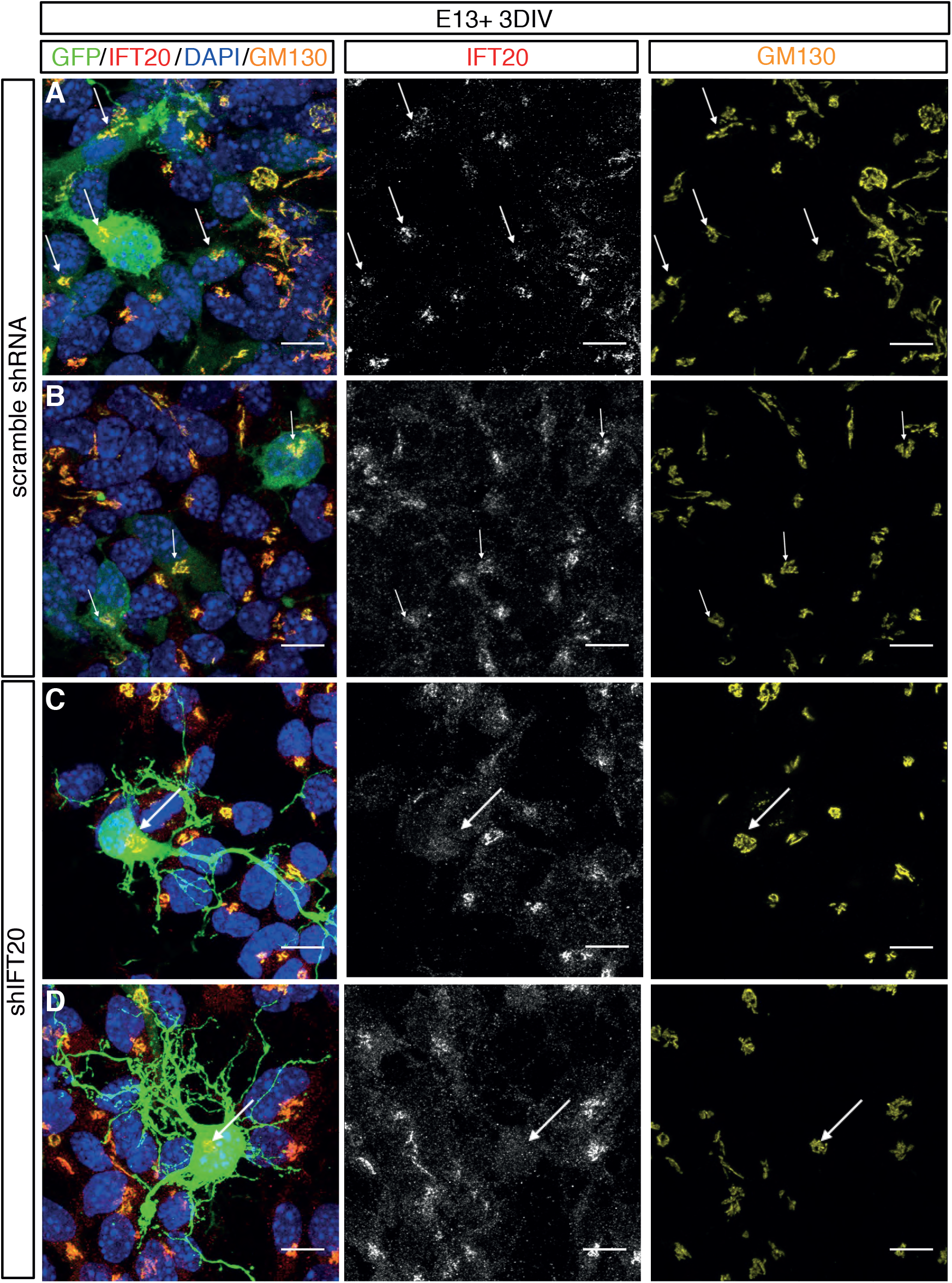
Validation of IFT20 shRNA in cortical cells in vitro. (A-D) Immunofluorescence staining of 2D mouse embryonic cortex primary culture with DAPI, IFT20 and GFP after ex utero electroporation performed at E13.5 and analysis 72H later. (A,B) scramble shRNA or (C,D) mix of sh targeting mouse IFT20 (shIFT20). Cells transfected with a mix of shRNA against IFT 20 show decrease signal for IFT20. (2 exp n= 4 microscopic fields/experiment). Scale bar 10 υm.

